# Topological Data Analysis of Protein Structure Manifolds from Molecular Dynamics Computer Simulation

**DOI:** 10.1101/2025.07.12.664527

**Authors:** Mirela Sino, Hiqmet Kamberaj

## Abstract

The analysis of computer simulation data requires efficient statistical and computational approaches, based on well-established theoretical frameworks. This study aims to introduce such approaches for topological data analysis within the persistent homology framework and to describe the manifold of the protein structure dynamics within the differential geometry of the directed graphs framework. Furthermore, the asymmetric kernel-directed graphs determined by the transfer entropy will describe the information flow in this manifold. The primary goal is to characterise changes in the topology of the protein structure due to the mutations. Moreover, this study aims to define the embedded manifold of dimension *m* of the amino acid sequence interaction network using the graph’s Laplacian matrix for determining the local embedded vector fields and coordinate vectors in this manifold for each amino acid as the vertices of either a directed or undirected graph.

Furthermore, this study strives to show that encoding the amino acid sequence information in an *m*-dimensional manifold is statistically efficient by decoding that information in a much lower-dimensional space. Then, using the topological data analysis, we can observe protein structure dynamics changes in a multidimensional manifold, for example, due to amino acid mutations. The analysis showed that short equilibrium structure fluctuations at a few nanoseconds enable the construction of such a manifold.

As a case study, the influence of the mutation of the two disulphide bridges on the three-dimensional structure of the Bovine Pancreatic Trypsin Inhibitor protein is investigated.

## 1 Introduction

Considerably large amounts of the data collected to investigate physical and chemical phenomena, such as the molecular dynamics (MD) simulations of biological molecular systems [54, 73, 78], may require efficient statistical and computational approaches. Furthermore, the cross-validation of these approaches requires well-established theoretical frameworks.

The problem of encoding the protein structure into amino acid sequences relates to the information contained in the amino acids, classified by an overlap of their physical and chemical properties. The information-theoretic measures, such as mutual information [51, 46] and transfer entropy [74, 79, 11, 58, 36, 57], have already been considered [46, 36] to be sensitive to linear and nonlinear statistical correlations. They can measure the statistical mutual information content between the amino acids of a biological system (considered as components of an interaction network system) by focusing on the position fluctuations of a group of atoms.

Furthermore, the mutual information or Pearson correlation coefficient are symmetric measures (that is, *R*_*ij*_ = *R*_*ji*_ for any two components *i* and *j* of an interaction network). Besides, the advances in computer architectures have made possible the acceleration of analysis and visualisation of large biological structures [40]. Thus, combining with advances in graph embedding and visualisation [50], it is possible to analyse the interaction networks of large amino acid sequences. However, the mutual information as a symmetric measure can identify the direction of the information flow; hence, it can not distinguish the *sender* (or *source*) and *receiver* (or *sink*) amino acids in an interaction network of the graph data [36]. Interestingly, the amino acid sequences are a class of interaction network graph data where the information content in the data is naturally asymmetric. This is supported by the experimental studies of the mutation analysis in the biological molecular systems [4, 5].

The transfer entropy is an asymmetric measure enabling measurement of the direction of the information flow in a directed graph data representation where the amino acids of a protein are the vertices of that graph [33]. The transfer entropy has been greatly appreciated in studying different physical, chemical, and biological phenomena in biological molecular systems [42, 36, 43, 48, 44, 2, 3, 33].

The statistical analysis framework is presented [33] in studying the amino acids interaction network associated with the graph topology of the information flow in proteins. The weighted directed graph model described mathematically the topological network of the anisotropic diffusion of the heat flow using the equilibrium structure fluctuations at sub-nanoseconds time scales from molecular dynamics simulations.

In a previous study [35], we cross-validated the statistical efficiency of the transfer entropy measures to determine measurements of high-order correlations that satisfy information-theoretic properties of information transfer. Furthermore, a machine learning approach was employed to classify the amino acids’ role in a protein structure.

This study aims to continue the previous efforts [35] to encode the amino acids’ information in an *m*-dimensional manifold by using statistically efficient theoretical frameworks. In particular, we aim to consider the asymmetry in data by studying the spectral properties of the directed Laplacian operator from a graph-theoretic viewpoint. Besides, the transfer entropy [21] was used as an information flow kernel to preserve the asymmetry information in the data and describe the connectivity.

This study introduces a topological data analysis within the persistent homology framework to examine the process of generating the directed graphs and the statistical properties of the operator. The primary goal is to characterise changes in the topology of the protein structure due to the mutations. Furthermore, the vertices of those directed graphs are viewed as a sub-nanoseconds sample from the manifold in an Euclidean space, and the edges represent the macroscopic measurements of the heat flow kernel between adjacent vertices on the manifold. We strive to determine the heat flow kernel upon the transfer entropy to characterise the overall directed graph connectivity and asymmetry, and the Laplacian operator generated from them explores the process of manifold sampling, concerns that are also investigated elsewhere [19]. Moreover, this study’s goal is to provide a theoretical framework determining the key features of the sampling process, such as the local embedded vector space and coordinate space of a manifold geometry, sampling distribution, and local information of the direction diffusion maps [19], useful for employing machine learning approaches.

This study will investigate the influence of disulphide bridges on the three-dimensional molecular structure of the Bovine Pancreatic Trypsin Inhibitor (BPTI) protein (in its wild and mutated types) [10, 38] using the data of molecular dynamics from our previous study [35].

## 2 Theoretical Frameworks

### 2.1 The (*α, q*) Transfer Entropy

The transfer entropy, *T*_*j*→*i*_, represents a dynamical measure, as a generalisation of the entropy rate (*H*) to more than one element to form a mutual information rate (*I*) [74], expressed as:

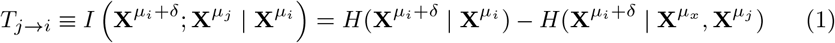

where 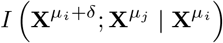 is the conditional mutual information [11, 36, 34]. Equation 1 indicates that the transfer entropy, *T*_*j*→*i*_, measures the average amount of information contained in the source (random process *j*) about next state 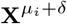 of the random process *i* that was not already contained in the past of process *i*.

Here, the time delayed embedding method is used to reconstruct the dynamics of the random time processes [33], *µ*_*i*_ = (*m*_*i*_, *τ*_*i*_). In particular, we construct the sets of state vectors 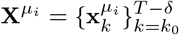 such that

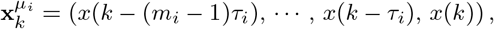

and 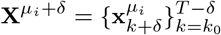.

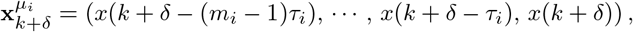

where *δ* is a time step representing the time length of influence. Taking *k*_0_ = max{(*m*_*i*_ − 1)*τ*_*i*_, (*m*_*j*_ − 1)*τ*_*j*_} creates vectors of the same length, *T* − *k*_0_ (where *T* is the length of the MD simulation run).

In our previous study [35], the statistical efficiency of alternative definitions of the transfer entropy measures between time series is validated, based on Sharma-Mittal (*α, q*)-framework [13], and the so-called (*α, q*) transfer entropy was defined as [35]:

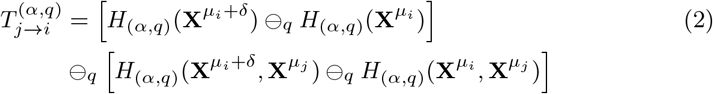

In Eq. (2), the function *x* ⊖_*q*_ *y* determines a modified difference as [37]:

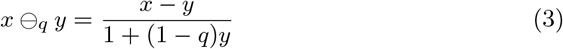

which reduces to standard difference for *q* = 1 (that is, *x*−*y*). In Eq. (2), *H*_(*α,q*)_(*X*) defines the Sharma-Mittal entropy [13] given in terms of Rényi entropy *H*_*α*_(*X*) [8, 68] as:

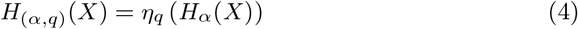

In Eq. (4), *η*_*q*_ is the following function [37]^1^:

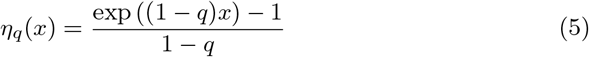

Note that for *q* = 1 and *α* ≠ 1, expressions reduce to Rényi transfer entropy formulations [68], based on *α*-framework of entropy while for *q* ≠ 1 and *α* = 1, they reduce to the so-called here Tsallis transfer entropies [14, 15], based on the so-called *q*-framework [35]. When *q* = *α* = 1, the standard transfer entropies based in the Shannon entropy formulation [74] are obtained as in Eq. (1), already used by one of us in the previous studies [36, 21, 33, 34].

It is worth emphasising that transfer entropy, *T*_*j*→*i*_, is sensitive to the non-linear causal correlations [36]; however, higher-order non-linear measures, as the ones described here 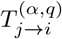, are more accurate in measuring weaker causal interaction correlations, such as dipole-dipole interactions.

### 2.2 Topology Manifold and Embedded Manifold

In the following, we are going to introduce some standard mathematical formulations in literature [7, 27, 64] for a general reader in structural molecular biology. It is worth noting that, to the best of our knowledge, some of these methodological frameworks have already been appreciated in the computational biology [30, 45, 6, 76, 81] and structural phase transitions in classical mechanics [52], including different areas of science [28, 29, 22].

An *m*-dimensional topology manifold ℳ is a topological space locally similar to Euclidean *m*-dimensional space ℛ^*m*^.

An embedded manifold ℳ is an *m*-dimensional space embedded in a higher-dimensional space ℛ^*n*^ (*m < n*). An embedded manifold always exists within a higher-dimensional space; however, a topological manifold does not necessarily exist within a higher-dimensional space. Furthermore, the computations are often carried out more easily in a topological space than in an embedded one. For example, molecular dynamics simulations are more conveniently carried out in the Cartesian coordinate space of the atoms than in an embedded topology, obtained by special techniques [60, 59, 32].

This study strives to show that the directional information flow kernel embedding DiGraphs can be used to describe the diffusion flow dynamics in an embedded manifold ℳ obtained using transfer entropy, considering that the entropic forces drive the flow over the manifold. It is worth noticing that the generative models for directed graphs using an asymmetric kernel matrix for weighted edges modelled from some vector field have already been discussed [19, 69, 20], using artificial and real data. Our algorithm presented in this study is essentially different because it is based on the transfer entropy, which is a causality measure that preserves the asymmetry of information flow contained in data.

The model consists of the observed DiGraph *G* of *m* vertices with affinity matrix **K**, as described below. Furthermore, assuming that *G* is a geometric random object composed of *m* vertices and using the Laplacian-kernel matrix **L**^(*t*)^ =(**X**^(*t*)^) ^*T*^ **X**^(*t*)^ approach, we sample *m* points according to the distribution *P* (**L**^(*t*)^) of the unobserved compact manifold ℳ in ℛ^*n*^ of some known intrinsic dimension *m ≤ n*. The directional attribute of the kernel from a vector field in ℳ determines the preferred direction between the DiGraph *G* weights *W*_*uv*_ and *W*_*vu*_, which is chosen here in terms of the information flow 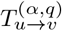 and 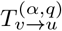, respectively. Moreover, the information flow can characterise any directionality related to the drift component of ℳ in ℛ^*m*^ and the spectral decomposition of the Laplacian matrix determines the embedding coordinate vectors **X**^(*t*)^ characterising the source components of local coordinates on the chart (*U*_*t*_, **x**^(*t*)^). While the transition probability matrix **P** characterises the random walk in a DiGraph and the drift component on ℳ (see also Refs. [19, 69, 20]).

Here, the topology of the data is characterised by the so-called affinity matrix (asymmetric similarity kernel) **K** with components *K*_*ij*_ measuring the strength of influence of *j* on *i*. The similarity kernel can be decomposed into two terms, namely the symmetric part **H** and anti-symmetric part **A**:

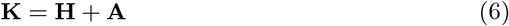

Where

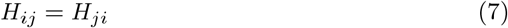

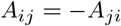

Here, we propose the transpose of the transfer entropy matrix as a newly introduced measure of asymmetric similarity kernel:

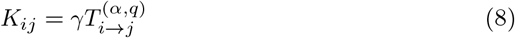

Where

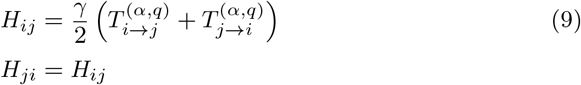

and

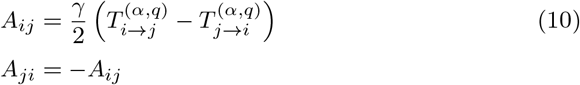

where *γ* is a real-valued constant, which defines the entire family of symmetric and asymmetric similarity kernels. Note that 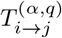 measures information flow from *i* to *j*; that is, in a DiGraph, *i* points (*visits*) to *j* (strength of influence of *j* on *i*) [34].

If the matrix **K** is used as the adjacency matrix of a DiGraph, then the local embedding tangent space vectors **V**^(*t*)^ are obtained at any time *t*. To obtain the local coordinates on the chart (*U*_*t*_, **x**^(*t*)^), we introduce a symmetric similarity kernel **K**^(*s*)^, as follows [19, 20]:

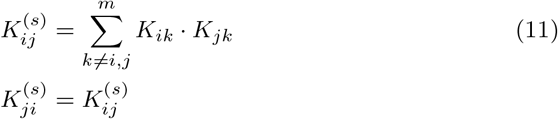

### 2.3 Affinity Kernel (Di)Graph Embedding

#### 2.3.1 DiGraph Embedding

Consider a directed graph *G*(*V, E*, **K**) with *m* vertices, *V* = {*v*_1_, *v*_2_, …, *v*_*m*_}, and *n* edges, *E* = {*ℓ*_1_, *ℓ*_2_, …, *ℓ*_*n*_} where *ℓ*_*i*_ = (*u, v*) is a directed edge from *u* to *v*, characterised an adjacency matrix **K**, which is the asymmetric kernel matrix defined above, with edges weights given as:

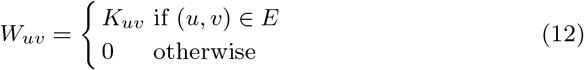

Next, two diagonal matrices can be determined to characterise the in- and out-degrees of vertices, denoted **D**^(in)^ and **D**^(out)^, respectively, as [35]:

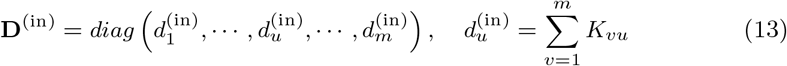

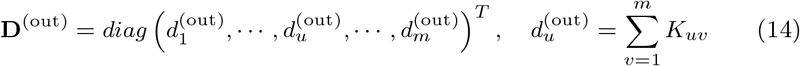

Then, the transition probability matrix 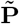 of rank *m × m* of a random walk on a DiGraph is given as [35]:

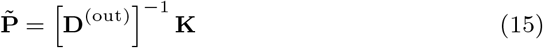

For a strongly connected digraph the random walk characterised by 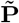 is an irreducible and aperiodic Markovian transition process, which is satisfied if there are no vertices with 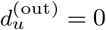 [33]. The following transition probability matrix is suggested to guarantee the irreducibility and aperiodicity [35]

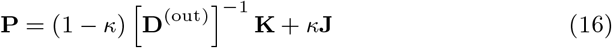

where **J** is a jumping matrix given in terms of the unitary diagonal matrix **1**_*m*_ of rank *m × m* as:

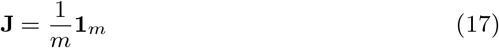

Here, *κ* is a number between 0.1 and 0.2 [33] (and the references therein). Then, the normalised Laplacian matrix is constructed as [31]:

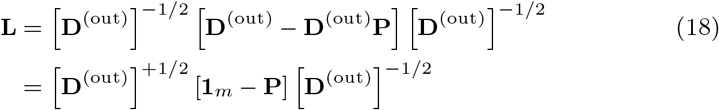

Spectral decomposition of **L** matrix is written as [70]:

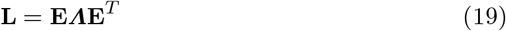

where **E** is the matrix of ordered eigenvectors, **e**_*u*_ = (*E*_1*u*_, …, *E*_*mu*_)^*T*^ as columns and ***Λ*** is a diagonal matrix of ordered eigenvalues (*λ*_1_ *< λ*_2_ *<* …*< λ*_*m*_), which are all non-negative. If we consider that the intrinsic dimension of the embedded vector space is *d « m*, then the embedding of *G* into ℛ^*d*^ determines the vector field components of a tangent space vector 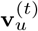 locally for the vertex *u* of *d* dimensions at any time *t* (for *u* = 1, 2, …, *m*, where *m* is number of sample points of the embedded manifold) as:

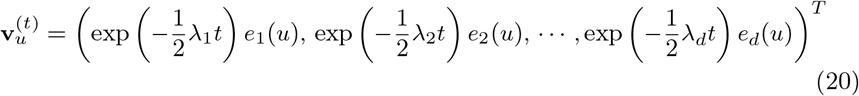

where *t* is the random walk diffusive time of directional flow on the directed graph and (^*T*^) stands for the transpose. Here, 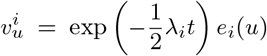 are the contravariant components of 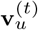 at time *t* for the vertex *u* in a local coordinate chart (*U*_*t*_, **x**^(*t*)^). In matrix form, **V**^(*t*)^ (a matrix with columns the vectors 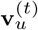):

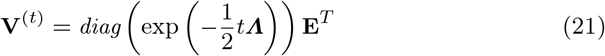

where *diag* stands for diagonal matrix with diagonal elements the components of vector 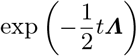 and off-diagonal elements equal to zero.

That represents a kernel local mapping at each vertex *u*:

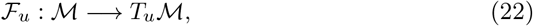

which embeds at each vertex of the DiGraph a tangent space *T*_*u*_ℳ.

#### 2.3.2 Graph Embedding

Now, consider a simple graph *G*(*V, E*, **K**^(*s*)^) with *m* vertices, *V* = {*v*_1_, *v*_2_, …, *v*_*m*_}, and *n* edges, *E* = {*ℓ*_1_, *ℓ*_2_, …, *ℓ*_*n*_} where *ℓ*_*i*_ = (*u, v*) is an edge between *u* and *v*, characterised an adjacency matrix **K**^(*s*)^, which is the symmetric kernel matrix defined above, with edges weights given as:

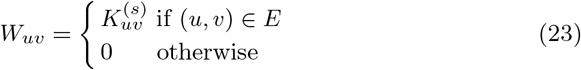

The normalised graph Laplacian is the matrix is defined as [31]:

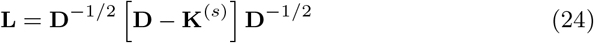

where **D** is the diagonal matrix of vertex degrees.

Similarly, spectral decomposition of **L** matrix gives [70]:

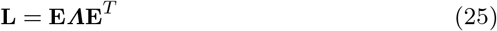

**E** is the matrix of ordered eigenvectors **e**_*u*_ = (*E*_1*u*_, …, *E*_*mu*_)^*T*^ as columns and ***Λ*** is a diagonal matrix of non-negative ordered eigenvalues (*λ*_1_ *< λ*_2_ *<* …*< λ*_*m*_). Again, considering the intrinsic dimension of the embedded vector space is *d « m*, then the embedding of *G* into ℛ^*d*^ determines the embedded local coordinate vector 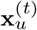 for the vertex *u* of *m* dimensions at any time *t* (for *u* = 1, 2, …, *m*) [31]:

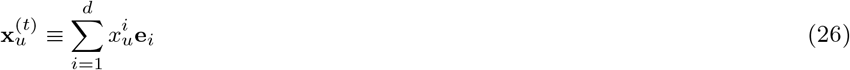

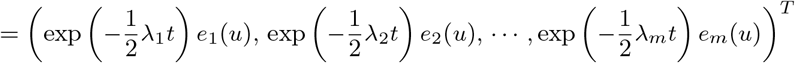

and in a matrix form, **X**^(*t*)^, with columns the vectors 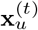:

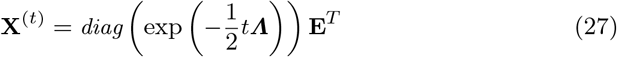

That represents a kernel mapping

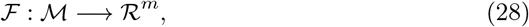

which embeds each vertex of the Graph into a coordinate space *R*^*m*^ of vectors. The distance between any two points **x** and **x** ^′^ in a tangent vector space equipped with a metric **g** is defined as:

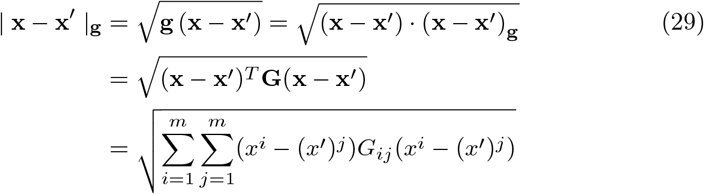

which is known as Mahalanobis distance [9] in a vector space. **G** is the matrix representation of the metric tensor **g**.

### 2.4 Topological Data Analysis

The topological data analysis (TDA) is a field dealing with the topology of the data to understand and analyse large and complex datasets [27, 30, 49, 41, 28].

#### 2.4.1 Persistent Homology

Persistent homology (PH) [27, 71] is part of TDA, aiming to construct a topological space gradually upon the input dataset, which is done by growing shapes based on the input data. PH is considered a topological tool for studying the non-linear global geometric data features. The features will be identified as *persistent* (and just a *noise*) if, after the last iteration, they are still present.

Persistent modules are the PH algebraic invariants of data types with topological spaces **T** and ℛ -valued functions *d* (such as the distances) *d* : **T** *→*ℛ. Consider **R** a set of real-valued numbers and **V**^(*k*)^ the vector spaces over a fixed field *k*. The persistent module, ℱ*H*_*i*_, is a function ℱ*H*_*i*_ : **R** *→* **V**^(*k*)^, where ℱ be a filtration on the dataset ℱ: **R** *→* **T**^(*k*)^, which maps an element of **R** to a topological space over a fixed field *k*. Here, ℱ is the sub-level set filtration 𝒮 (*d*) as:

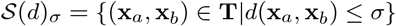

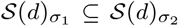 whenever *σ*_1_ *≤ σ*_2_; therefore, it is a filtration. The *i*th singular homology function *H*_*i*_ of some fixed field *k* is defined as

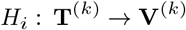

The persistence module ℱ*H*_*i*_, for every *i* ≥ 0, algebraically encodes the geometric information about the manifold of the dataset [71].

Let *M* be the persistence module with a finite-dimensional vector space. Then, there exists a unique way to decompose *M* into indecomposable interval persistence modules [77]. The intervals are characterised an indexing 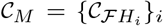 of the indecomposable collected intervals associated with *M*, the so-called barcodes of *M*, where *i* runs over the intervals or topological features. Here, 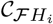 characterises the topological feature of an interval in the dataset, where the length of the interval is a measure of the significance of that feature.

#### 2.4.2 Persistance Diagram

Visualisation of the data point measurements is limited to two- or three-dimensional points in a chart. For higher-dimensional data points, the graphical representation becomes impossible. Furthermore, using the statistical summary (such as centre of mass, median, variance, and so on) may not be clear since they might be invariant under rotation, translation, and scaling of data. In contrast, the geometry of the topological space is preserved under these transformations.

Therefore, the graph theory provides an elegant and exact way to quantify geometrical shapes of discrete datasets. Typically, a graph *G* = (*V, E*) is a tuple of the set of *N* vertices *V* = {*v*_1_, *v*_2_, …, *v*_N_} and *K* edges *E* = {*e*_1_, *e*_2_, …, *v*_*K*_} where an edge *e*_*k*_ = (*u, v*) is said to connect the vertices *u* and *v* if they are sufficiently close. Then, to quantify topological invariants of the graph, one can compute Betti numbers *B*_*k*_, which represent an *k*-dimensional hole [17, 22]. Constructions of generalisations of graphs are known as simplicial complexes, which enable obtaining higher-dimensional objects, such as faces, volumes, and so on, through triangulation. Typically, a *k*-simplex is a combination of *k* + 1 vertices, and a *k*-simplicial complex is a set of simplices with at least *k* dimensions.

Persistence diagrams consist of points (*b, d*) (where *d* stands for *distraction* (or *death*) and *b* for *birth*) that represent different topological features. Typically, along the horizontal axis characterises the length scales *σ*_*b*_ at which a feature is created, and the vertical axis characterises a length scale *σ*_*d*_ at which that feature is destroyed. The distance from the dashed diagonal line indicates how long a feature persists over a length scale; that is, further a point from the diagonal dashed line, larger the persistence interval over the length scale is and it is said to have a longer *lifetime*, computed as *ℓ* = *d* − *b >* 0 (and hence, there are no points below the diagonal dashed line [22]).

### 2.5 Relationship Between Topology and Protein Structure Dynamics

The topology has played an important role in studying the phase space transitions of different types of systems [61, 62, 16, 52]. Therefore, the relationship between the topology and protein structure dynamics should give insights into the structure changes of a protein, for example, due to mutations. In Ref. [56], the topological and geometrical hypotheses in studying Hamiltonian dynamics of systems undergoing phase transitions in configurational space were described. Furthermore [52], the configurational phase space transitions should be associated with geometrical changes of the manifolds underlying Hamiltonian flows; moreover, these geometrical changes are associated with topological changes of the potential energy level sets in the protein’s configuration space.

The potential energy *U* (**r**_*g*_), with **r**_*g*_ = (**r**_1_, …, **r**_*g*_), level set is defined as a finite set of equipotential hyper-surfaces in an *g* degrees of freedom configuration space of the protein: *S*_*U*_ = {*u*_1_, *u*_2_, …*u*_*k*_} where 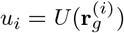 and *k* number of the levels. Here, *U* (**r**_*g*_) is a function *U* : ℳ *→* ℛ mapping a topological space ℳ to a real-valued set ℛ.

The manifolds bounded by *S*_*U*_ are defined as a set of ellipsoids 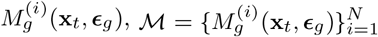. Here, 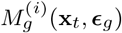 determines an ellipsoid of exes lengths ***E***_*g*_ and origin **x**_*t*_ at time *t*, which is a subspace element such that 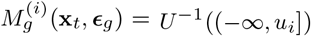. In fact, from this definition, *S*_*U*_ is a *g*-dimensional configurational space. Furthermore, the number of ellipsoids needed to cover the manifolds bound by *S*_*U*_ is *k ∼* (1*/E*^*g*^). The Hamiltonian flow in a high-dimensional space depends on the curvature of the hypersurface in the configuration space manifold [52].

It is worth noticing that compared to the topological data analysis, this represents a *filtration* of a simplicial complex *K*, which is a family of sub-complexes *K*_*i*_ (for *u*_*i*_ ∈ *V* and *V ⊆* ℛ). For any *u*_*i*_, *u*_*j*_ ∈ *V*, when *u*_*i*_ *≤ u*_*j*_, then *K*_*i*_ *⊆ K*_*j*_, and 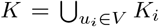. Furthermore, the filtration of the topological space ℳ is a set of subspaces *M* ^(*i*)^ with *u*_*i*_ ∈ *V* and *V ⊆* ℛ such that whenever *u*_*i*_ *≤ u*_*j*_, then *M*^(*i*)^ ⊆ *M*^(*j*)^, and 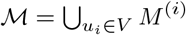. Hence, for a function *U* : ℳ → ℛ, the set of subspaces *M* ^(*i*)^ = *U*^−1^((−*∞, u*_*i*_]) with *u*_*i*_ ∈ ℛ determines the sub-level set filtration of *U*. The parameter *u*_*i*_ defines the *resolution* of the filtration.

The topological dependence of the structure changes rely on the relationship between the configurational entropy and topological properties of the manifolds 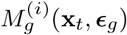 [56]: 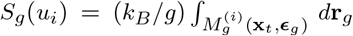 where *k*_*B*_ is the Boltzmann’s constant. Therefore, significant changes on the topology with *u*_*i*_ of the manifolds 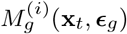 (which results in *sharp* changes of the potential energy function) can influence on the dependence of *S*_*g*_ (*u*_*i*_) (and its derivatives) on *u*_*i*_. As a result, it has been supposed that for some physical systems the configuration space transitions originates from changes on the topology manifolds 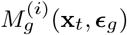 of the potential energy level set *S*_*u*_ [47, 66, 55, 53].

## 3 Molecular Dynamics Simulations of Modelled Systems

The initial protein structure for molecular dynamics simulations is the BPTI protein with PDB ID 5PTI, a protein of 58 amino acids, obtained by joint X-ray and neutron experiments at a resolution of 1.00 Å [10]. The structure contains three disulphide bonds, namely Cys5-Cys55, Cys14-Cys38, and Cys30-Cys51. For BPTI[14-38]Abu mutant (which has only one disulphide bond, namely Cys14-Cys38, and the other cystine amino acids Cys5, Cys30, Cys51, Cys55 are mutated to *α*-amino *n*-butyric acid), the three-dimensional initial structure was modelled as in Ref. [35] from the X-ray experiments at a resolution of 1.70 Å [38] (PDB ID 1YKT). The molecular dynamics simulation details and data acquisition are described elsewhere [35].

## 4 Results and Discussion

### 4.1 Structural Stability Analyses

#### 4.1.1 Long-range Electrostatic Interactions

Figure 1 presents the trajectories of the salt-bridges (as O .. N distances in angstrom) in the wild-type BPTI (BPTI-WT) and [14-38]Abu with Ala11Thr, Arg15Lys, Lys26Pro, and Ala27Asp mutations (BPTI[14-38]Abu) during the last

**Figure 1.**
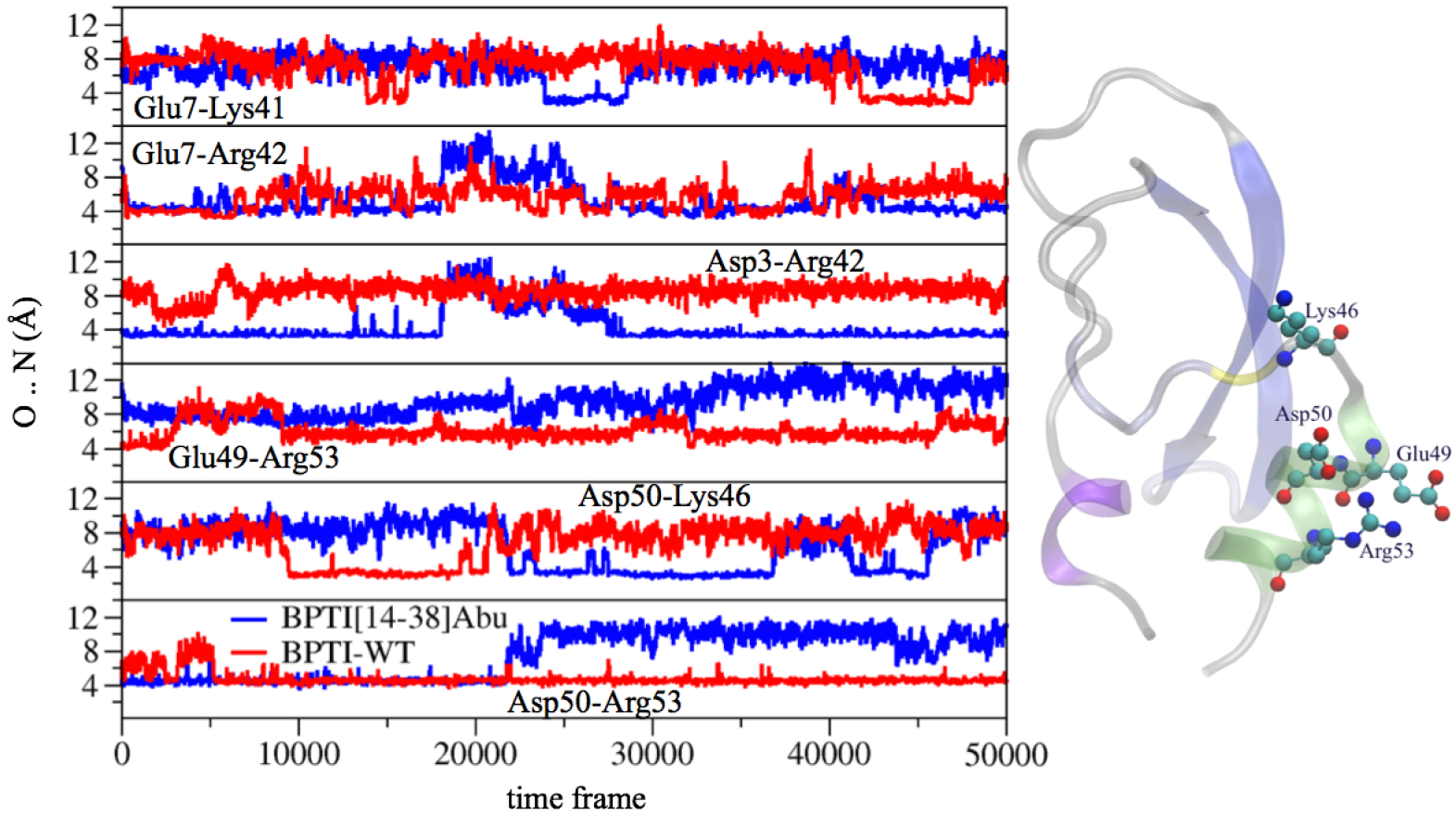
The O .. N distance trajectories of the wild-type BPTI and [14-38]Abu with Ala11Thr, Arg15Lys, Lys26Pro, and Ala27Asp mutations. In the last 10 ns of MD simulation runs, the configurations were saved and analysed every 0.2 ps.

10 ns of MD simulation runs. The configurations were saved every 0.2 ps. These analyses help to understand the influence of long-range electrostatic interactions on the structural stability. Our results show a destabilisation of Glu49−Arg53 and Asp50−Arg53 long-range electrostatic interactions and a strengthening of Asp50−Lys46 interaction for BPTI[14-38]Abu mutant compared to BPTI wild-type. This may increase the overall stability of BPTI[14-38]Abu protein tertiary structure with the cost of decreasing the stability of secondary helical structure, which is influenced by Glu49−Arg53 and Asp50−Arg53 salt-bridges, as depicted in Fig. 1.

#### 4.1.2 Structural Fluctuations

Figure 2 shows the root mean square fluctuation (rmsf) of an amino acid in the wild-type BPTI-WT and BPTI[14-38]Abu mutant, based on C_*α*_ atom positions (where most of the fluctuations for the large-scale motions are concentrated [1]). The last 10 ns of MD simulation runs were used to analyse the results, saved every 2 ps, after the overall translational and rotational motions were removed [63, 72]. The residues indexed between 1 and 8 show higher fluctuations in BPTI[14-38]Abu structure compared to the BPTI-WT one, indicating a decrease in stability of the 3_10_-helix secondary structure. Similarly, the *α*-helix (residues indexed between 50 and 57) becomes unstable in the structure of BPTI[14-38]Abu (as depicted from Fig. 2).

**Figure 2.**
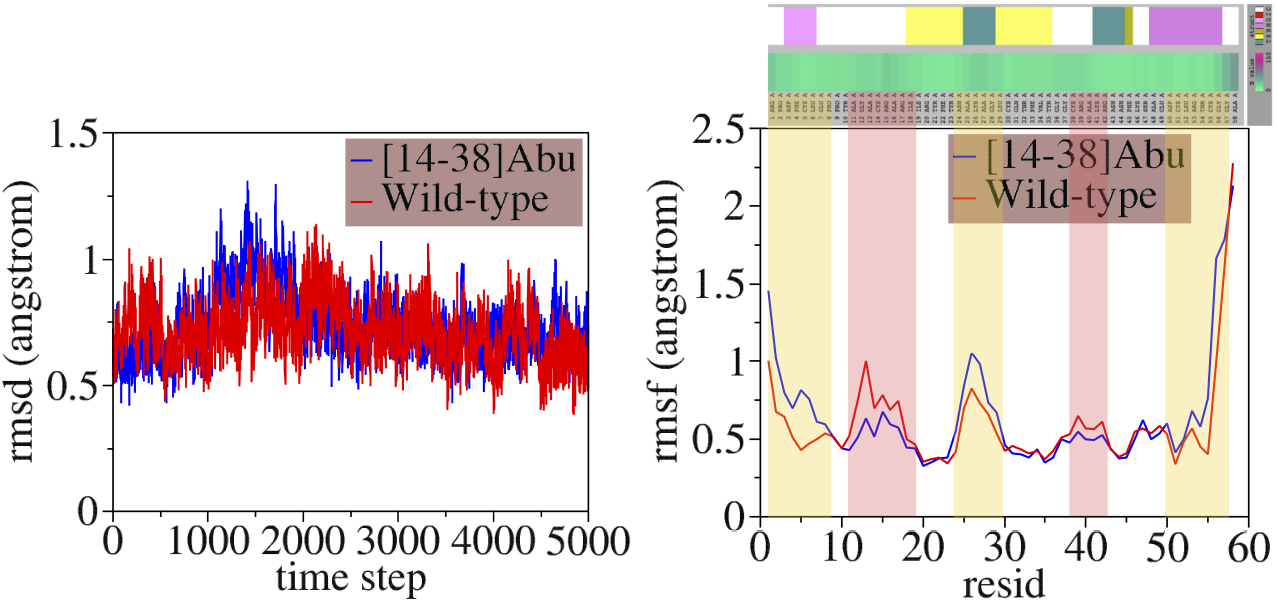
The rmsd and rmsf of the wild-type BPTI-WT and BPTI[14-38]Abu mutant. The last 10 ns of MD simulation runs were used to analyse the results, and the configurations were saved every two ps.

Furthermore, the root mean square fluctuation analyses indicate a decrease in stability of the residues 24-29 forming the turn-like secondary structure that links the two *β* sheets (Fig. 2). Interestingly, the coil residues (with ID between 11 and 18) have less fluctuations in BPTI[14-38]Abu mutant compared to wild-type BPTI-WT, indicating an increase in the stability of the first *β*-sheet secondary structure. The same increase in stability is observed for the turn structure with residues indexed from 38 to 42 in the BPTI[14-38]Abu protein.

### 4.2 NMR - Nuclear Magnetic Resonance

Next, the nuclear Overhauser enhancement (NOE) was computed to determine the spin-lattice dipolar relaxation contributions [25], using the CHARMM program [12] implementation. Figure 3 presents the spin-spin NH−HN sequential amino acid NOEs for the BPTI-WT (in blue) and BPTI[14-38]Abu (in red). Small values of NOEs imply high flexibility. The most interesting is the role of residues Ala16 and Ala58 (see also Fig. 3). That is, in the BPTI[14-38]Abu protein, the NOE of Ala16 is positive, and its NOE in wild-type protein is negative; in contrast, in the BPTI[14-38]Abu structure, the NOE of Ala58 is negative, and it changes to a positive value in the wild-type protein structure, BPTI-WT. That explained the increase in the intensity of medium-range NOE of Ala16 in BPTI[14-38]Abu mutant, which was confirmed by computing NH−NH non-sequential Ala16 NOEs with other amino acids. From our results, the Ala16N−Cys38N proton pair was the most intense NOE of around 1.077, which is two residues of the coil region linking the two *β*-sheets. This proton pair’s so-called effective relaxation time was around 236.89 ps in the BPTI[14-38]Abu structure. It is worth noting that the distance between the pair Ala16N−Cys38N, based on X-ray structure, is 6.83 Å [38] in BPTI[14-38]Abu. These data indicate that in BPTI[14-38]Abu, the flexibility of residues 16-19 reduces, hence increasing the stability of the *β*_1_-sheet.

**Figure 3.**
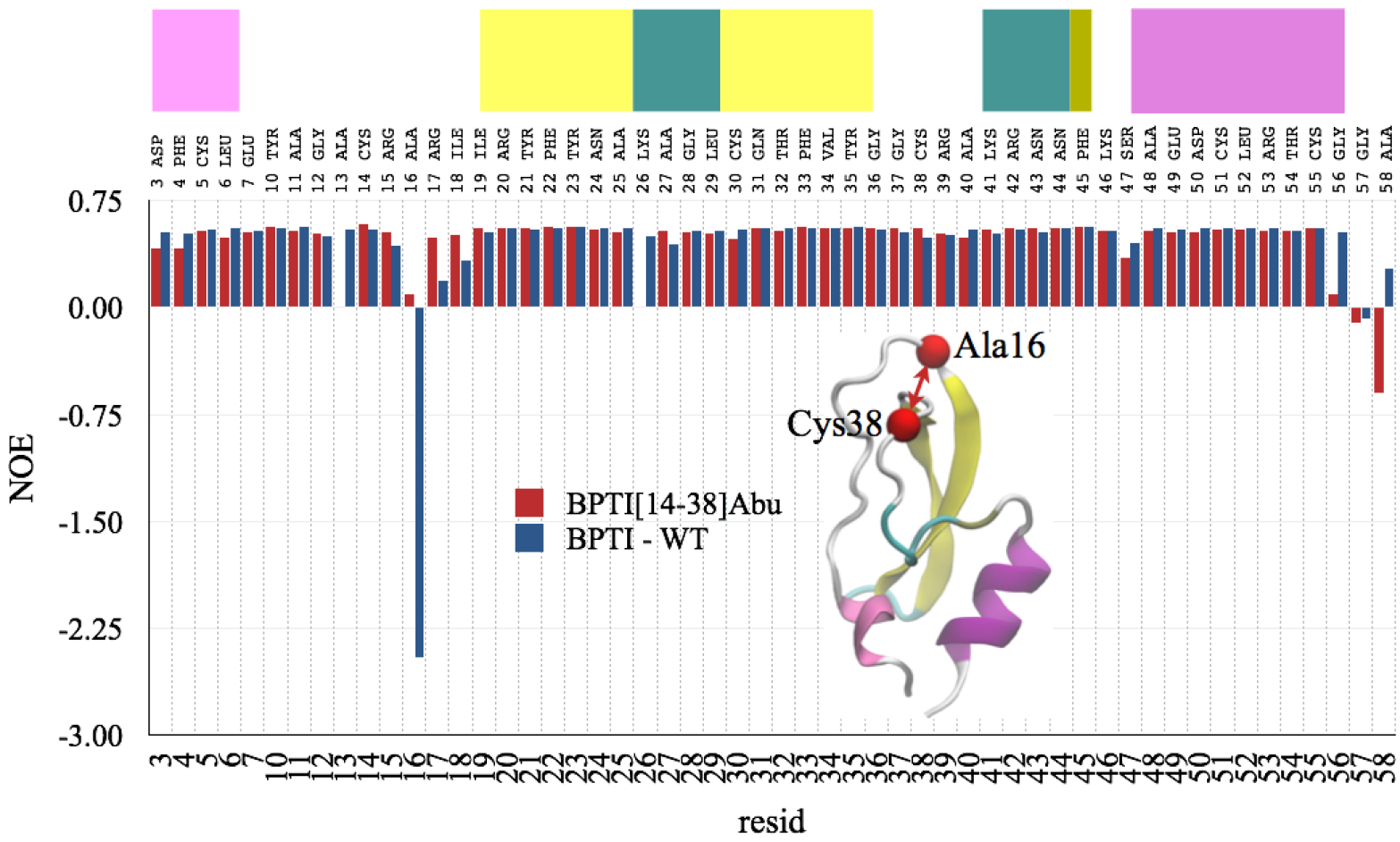
The NOE of the wild-type BPTI-WT and BPTI[14-38]Abu obtained by the last 10 ns MD simulation runs with configurations saved every 2 ps. The cutoff distance of the atoms pair was 2.0 Å, magnetic field was 11.74 Tesla, and the overall rotational tumbling motion period was 3000 ps.

Furthermore, Tab. 1 shows the NMR spectra (see also Ref. [67]). The CHARMM program [12] was used to compute the spectra for the turn region, namely the residues with ID between 25-28, using the medium-range proton pairs N−N. The equilibrium distances *d* are according to crystal structure experiments from references [10, 38]. Our results indicate that the NOE are more vital for the wild-type BPTI-WT than BPTI[14-38]Abu mutant. These results support experimental findings [23] that residues 25-28 stabilise turn conformations that nucleate folding, which may prevent mis-folding of *β*_1_ and *β*_2_ sheets.

Moreover, the N−HN dipole-dipole relaxation correlation functions are computed. Figure 4 shows the estimated effective correlation time *τ*_*e*_ (in ps) for the NH bond motions of the amino acids. Clearly, the results indicate the influence of mutations on the NH motion relaxations. In particular, the *α*-helical structure (including residues 46-54) of the wild-type (BPTI-WT) is characterised by a faster relaxation time compared to the mutated analog (BPTI[14-38]Abu). Similar results are obtained for Ala16 and residues 18-20 of coil region connecting *β*-sheet (*β*_1_), as depicted from Fig. 4.

**Figure 4.**
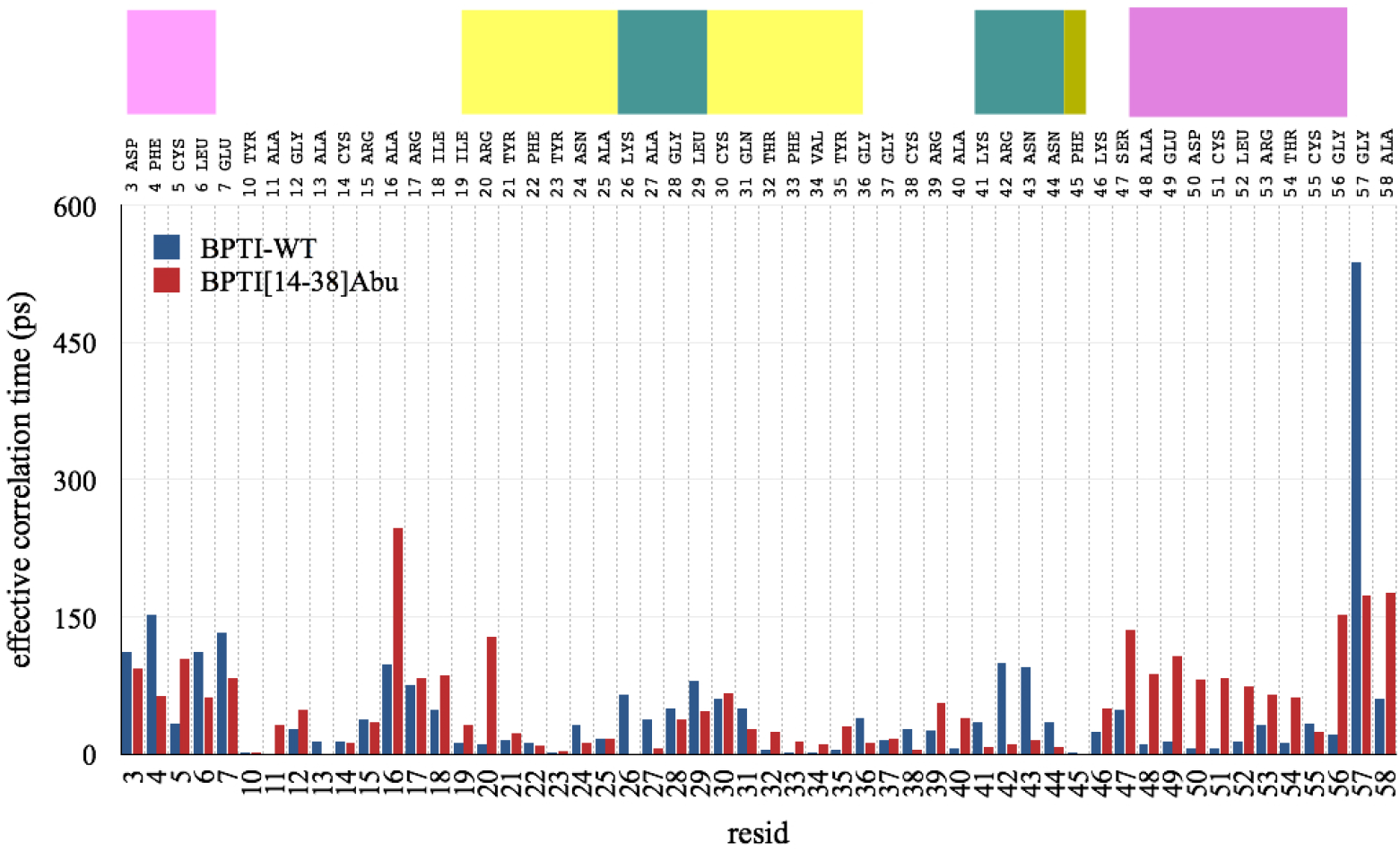
The effective correlation times *τ*_*e*_ (in ps) for the wild-type BPTI-WT and BPTI[14-38]Abu mutant, as obtained from the N−HN dipole-dipole relaxation correlations functions.

Table 2 presents the NMR spectra for the 3_10_-helix secondary structure, residues indexed between 3-6, using the medium-range proton pairs N−N. The NH bonds cross peaks are sharper in the BPTI[14-38]Abu protein structure than wild-type BPTI-WT one, indicating that the proton pairs are not in an intermediate exchange time regime and do not sample multiple conformations. Interestingly, these findings agree with NMR experimental results [23, 80]. They are also supported by the observation of the effective correlation times, presented in Fig. 4, which show lower effective correlation relaxation times for BPTI[14-38]Abu compared to BPTI-WT. It is worth noting that TMXE denotes the first time when the correlation function *C*(*t*) is less than or equal to the mean bond order parameter *(S*_2_), measured in picoseconds [65, 12].

**Table 1:** The NOE, effective relaxation time *τ*_*e*_, and TMXE (which is the first time when the time correlation *C*(*t*) is less than or equal to the order parameter of the spin-spin bond), according to Ref. [67] and computed using the CHARMM program [12], for the residues with ID between 25-28 using the medium-range proton pairs N−N. The equilibrium distances *d* are from crystal structure experiments [10, 38].

| Proton pair | NOE | $\tau_e$ [ps] | TMXE [ps] | $d$ [Å] |
| --- | --- | --- | --- | --- |
| BPTI-WT |  |  |  |  |
| Ala25N-Lys26N | 1.079 | 222.00 | 2094.0 | 2.72 |
| Ala25N-Ala27N | 1.078 | 136.00 | 1546.0 | 4.38 |
| Ala25N-Gly28N | 1.077 | 12.73 | 194.0 | 4.69 |
| Lys26N-Ala27N | 1.077 | 9.27 | 159.0 | 2.83 |
| Lys26N-Gly28N | 1.075 | 12.00 | 154.0 | 4.24 |
| Ala27N-Gly28N | 1.076 | 10.89 | 163.0 | 2.70 |
| BPTI[14-38]Abu |  |  |  |  |
| Ala25N-Lys26N | 1.075 | 24.49 | 211.0 | 3.04 |
| Ala25N-Ala27N | 1.071 | 36.29 | 279.0 | 4.77 |
| Ala25N-Gly28N | 1.075 | 44.59 | 439.0 | 4.55 |
| Lys26N-Ala27N | 1.074 | 42.33 | 376.0 | 3.01 |
| Lys26N-Gly28N | 1.077 | 51.49 | 474.0 | 3.99 |
| Ala27N-Gly28N | 1.076 | 39.12 | 280.0 | 2.62 |

**Table 2:** The NOE, effective relaxation time *τ*_*e*_ (in ps), and TMXE (in ps), according to Ref. [67] and computed using CHARMM program [12], for the 3_10_-helix region (residues 3-6), using the medium-range proton pairs N−N. The equilibrium distances *d* are obtained from data in Refs. [10, 38].

| Proton pair | NOE | $\tau_e$ [ps] | TMXE [ps] | $d$ [Å] |
| --- | --- | --- | --- | --- |
| BPTI-WT |  |  |  |  |
| Asp3N-Phe4N | 1.079 | 297.5 | 2162.0 | 2.83 |
| Asp3N-Cys5N | 1.078 | 109.0 | 1068.0 | 4.41 |
| Asp3N-Leu6N | 1.075 | 83.4 | 639.0 | 5.60 |
| Phe4N-Cys5N | 1.078 | 238.0 | 2246.0 | 2.72 |
| Phe4N-Leu6N | 1.077 | 132.9 | 1687.0 | 4.38 |
| Cys5-Leu6N | 1.076 | 55.2 | 949.0 | 2.71 |
| BPTI[14-38]Abu |  |  |  |  |
| Asp3N-Phe4N | 1.086 | 233.2 | 1032.0 | 2.85 |
| Asp3N-Abu5N | 1.082 | 182.5 | 996.0 | 4.29 |
| Asp3N-Leu6N | 1.072 | 83.6 | 304.0 | 5.42 |
| Phe4N-Abu5N | 1.084 | 351.2 | 2049.0 | 2.71 |
| Phe4N-Leu6N | 1.074 | 79.3 | 347.0 | 4.34 |
| Abu5-Leu6N | 1.072 | 71.6 | 349.0 | 2.67 |

The slow and fast relaxation time rates (in s^−1^) are computed for BPTI-WT and BPTI[14-38]Abu to further support our findings presented above. They are shown graphically in Fig. 5. A small value indicates a long relaxation time, and a large one indicates a short relaxation time of the amino acid in the protein structure. Our results (see Fig. 5) show that the slow relaxation time rates are in the range from 1.0 to 2.5 (s^−1^) and fast ones are between 1.8 and 4.5 (s^−1^), in agreement with previously reported experimental observations [24]. Similarly to the above-reported findings, such as Ala16, the slow and fast relaxation time rates of BPTI-WT are shorter than BPTI[14-38]Abu; that is, Ala16 is more flexible in the wild-type compared with the mutant analog.

**Figure 5.**
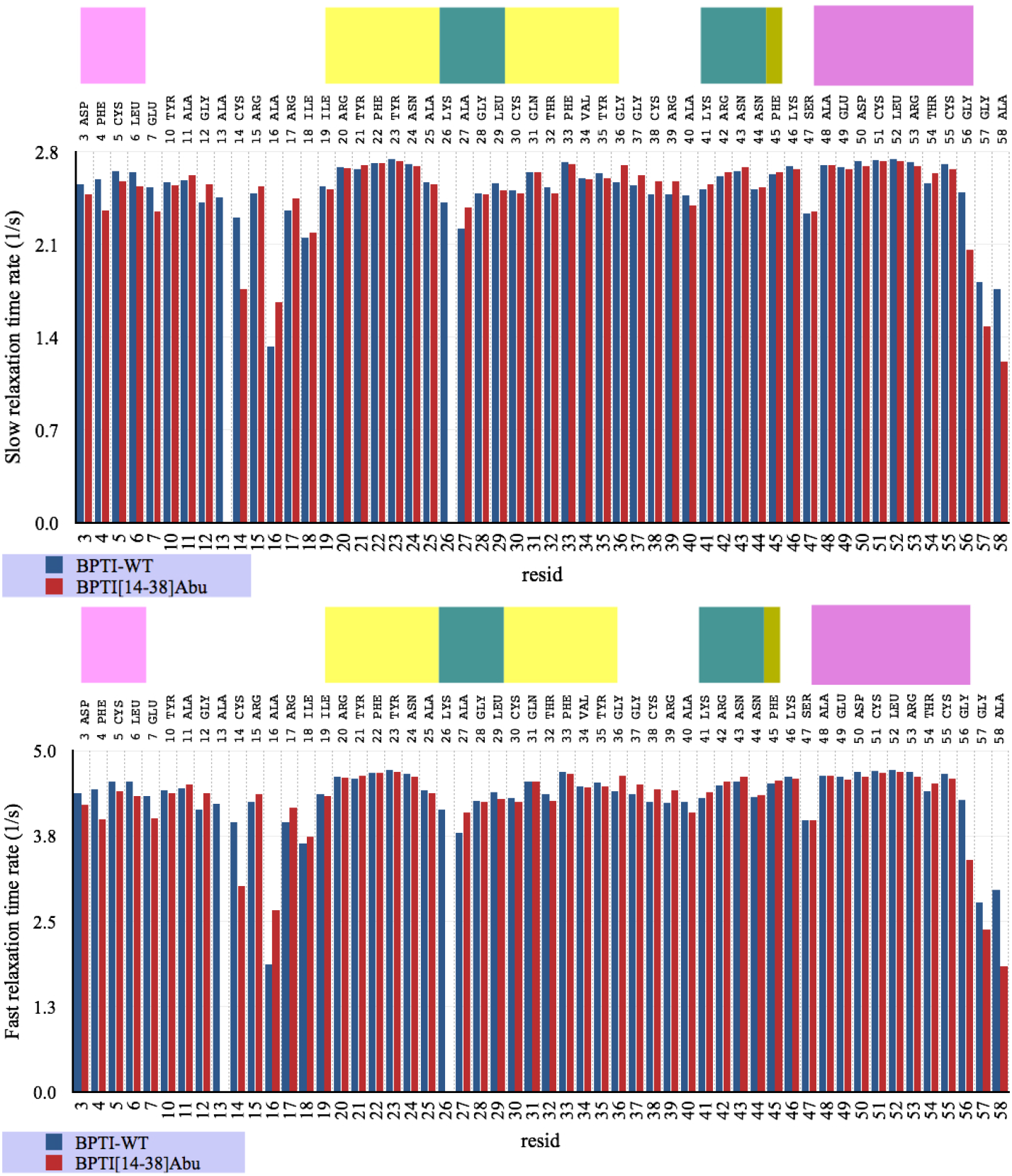
The slow and fast relaxation time rates (in s^−1^) for the wild-type BPTI-WT and BPTI[14-38]Abu structures from the last 10 ns MD simulation runs, computed every 2 ps.

Similar results are also confirmed by the observation of N…HN vector order parameter, ⟨*S*_2_⟩, which is a plateau value of the generalised order parameter [39], computed from an ensemble average over 5000 configurations saved from the last MD simulation runs [12]. Fig. 6 shows ⟨*S*_2_⟩ for each amino acid in the wild-type BPTI-WT and BPTI[14-38]Abu structure. A large value of ⟨*S*_2_⟩ represents a more rigid vector, and a small value indicates a more flexible bond vector. Clearly, our results confirm the increase of the rigidity of the N−HN bond vector for Ala16 in the BPTI[14-38]Abu mutant compared to BPTI-WT wild-type. In contrast, for Gly56 and Ala58, these vector bonds are more aligned in the wild-type BPTI-WT.

**Figure 6.**
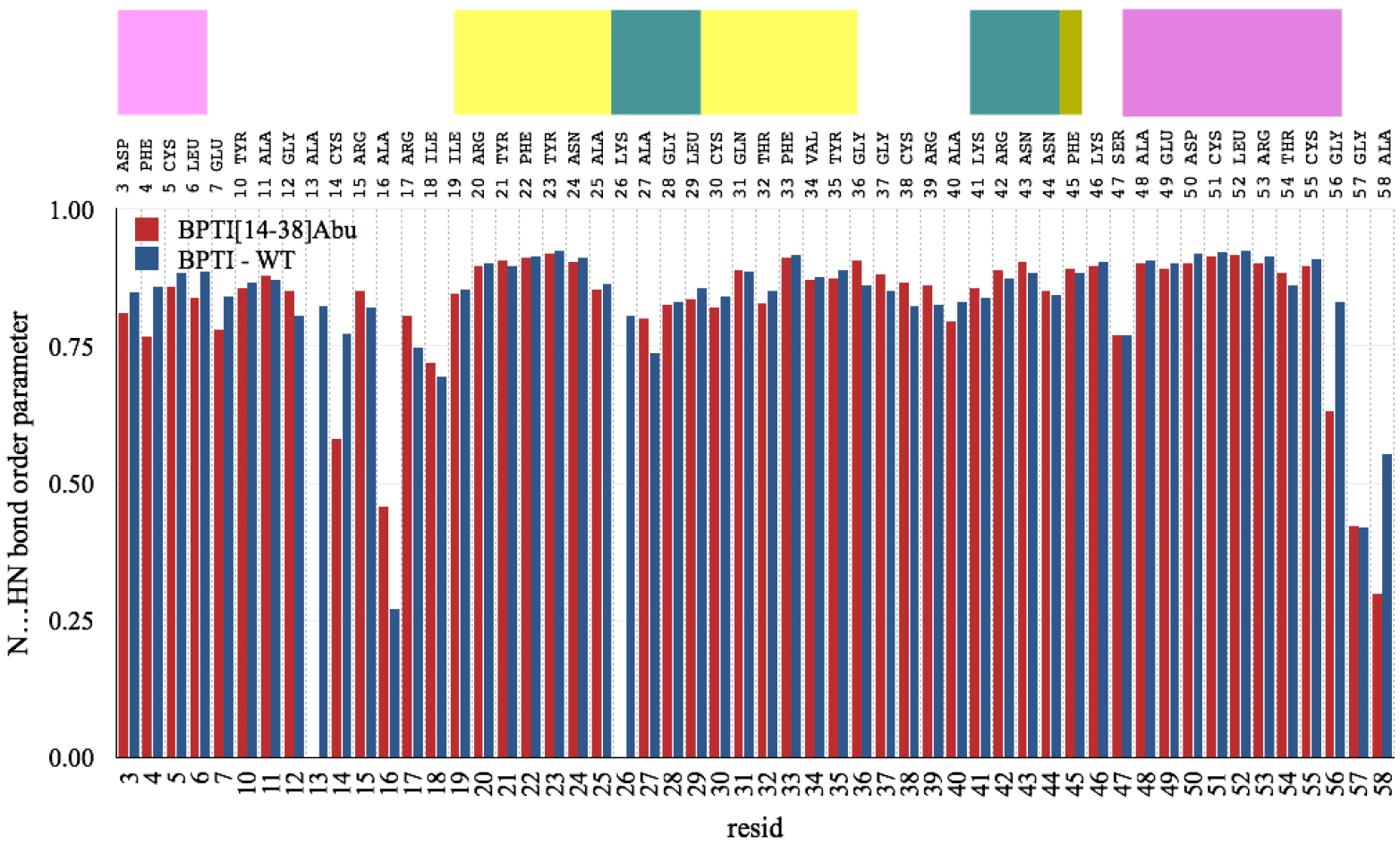
The order parameter of N…HN bond vectors for the wild-type BPTI-WT and BPTI[14-38]Abu structures, computed from the plateau of the generalised order parameter [39].

### 4.3 Dipole-dipole Interactions Information Flow

The so-called *directional information flow*, 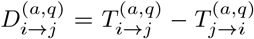 [36], calculated using the Sharma-Mittal symbolic encoding transfer entropies are shown in Fig. 7. The choice was *α* = 2.0 and *q* = 0.85 [35]. Here, if 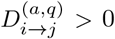, then information flows from source *i* to target *j*, or *i* influences *j*. Figure 7 presents heat map plots of 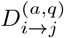 (in bits) for BPTI-WT and BPTI[14-38]Abu, where a red colour indicates a strong *source* of information and a blue one indicates a *target* (or *sink*) amino acid of information flow. Our results clearly identify the source and target residues and the change in the amino acid’s propensity to be a *source* or *target* (*sink*) of the information flow matrices for BPTI-WT and BPTI[14-38]Abu.

**Figure 7.**
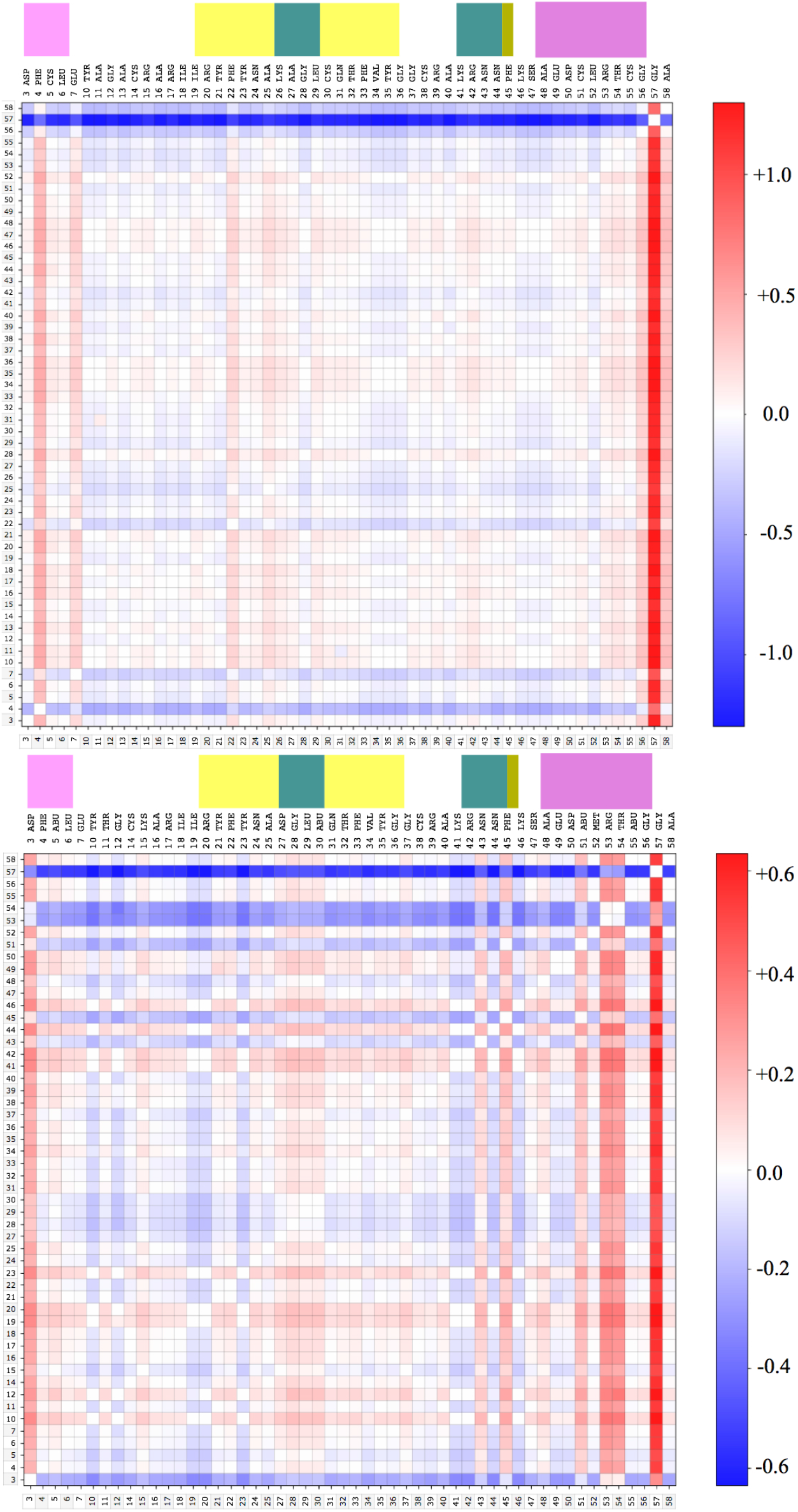
The comparison of directional information flow 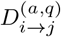 (in bits) for BPTI wild-type and BPTI[14-38]Abu.

Our results (see also Fig. 7) show that the residues 4-7 of the 3_10_-helix region change from target in the BPTI-WT structure to source of dipole-dipole interaction information flow in the BPTI[14-38]Abu. This is due to the strengthening of dipole-dipole interactions between 3_10_-helix and the turn region (residues 27-30) linking *β*_1_- and *β*_2_-sheets in the BPTI[14-38]Abu compared to wild-type BPTI-WT. Furthermore, those residues of 3_10_-helix becomes sources of information flow concerning the interactions with residues 43, 45 (of the turn linking *β*_2_ sheet with extended *β*-sheet), 48, 51, 53, 54, and 57 (of *α*-helix).

### 4.4 Topological Data Analysis

Figure 8 shows the topologies of the manifolds of BPTI wild-type and BPTI[14-38]Abu based on the dipole-dipole directed graph distance matrix. Here, we use the shortest path distances between nodes to construct a so-called *metric space*. The length of a path can be given either as the number of edges (*L*_0_) or as the total length of its edges (*L*_1_, which takes into consideration the edges’ weights). Importantly, if the graph is directed, then it is possible that *not all* nodes are reachable by a directed path from every other node (not strongly connected). In this case, we assigned the ∞ value to be the distance between unreachable pairs of nodes. In Fig. 8, *H*_0_ denotes the isolated points algebraic topology (connected components), *H*_1_ line segments (or holes), *H*_2_ triangles (or cavities), *H*^*i*≥3^ poly-edges shapes (non-cavities); for example, *H*_3_ corresponds to a tetrahedron algebraic topology shape. Our results indicate different elements of the topological space structures for BPTI-WT and BPTI[14-38]Abu; in particular, the cavities and *H*_3_ (poly-edges) elements of the topological space sets appear more often in BPTI[14-38]Abu than BPTI-WT. These results show that the topological space of BPTI[14-38]Abu is more complex than the BPTI-WT one.

**Figure 8.**
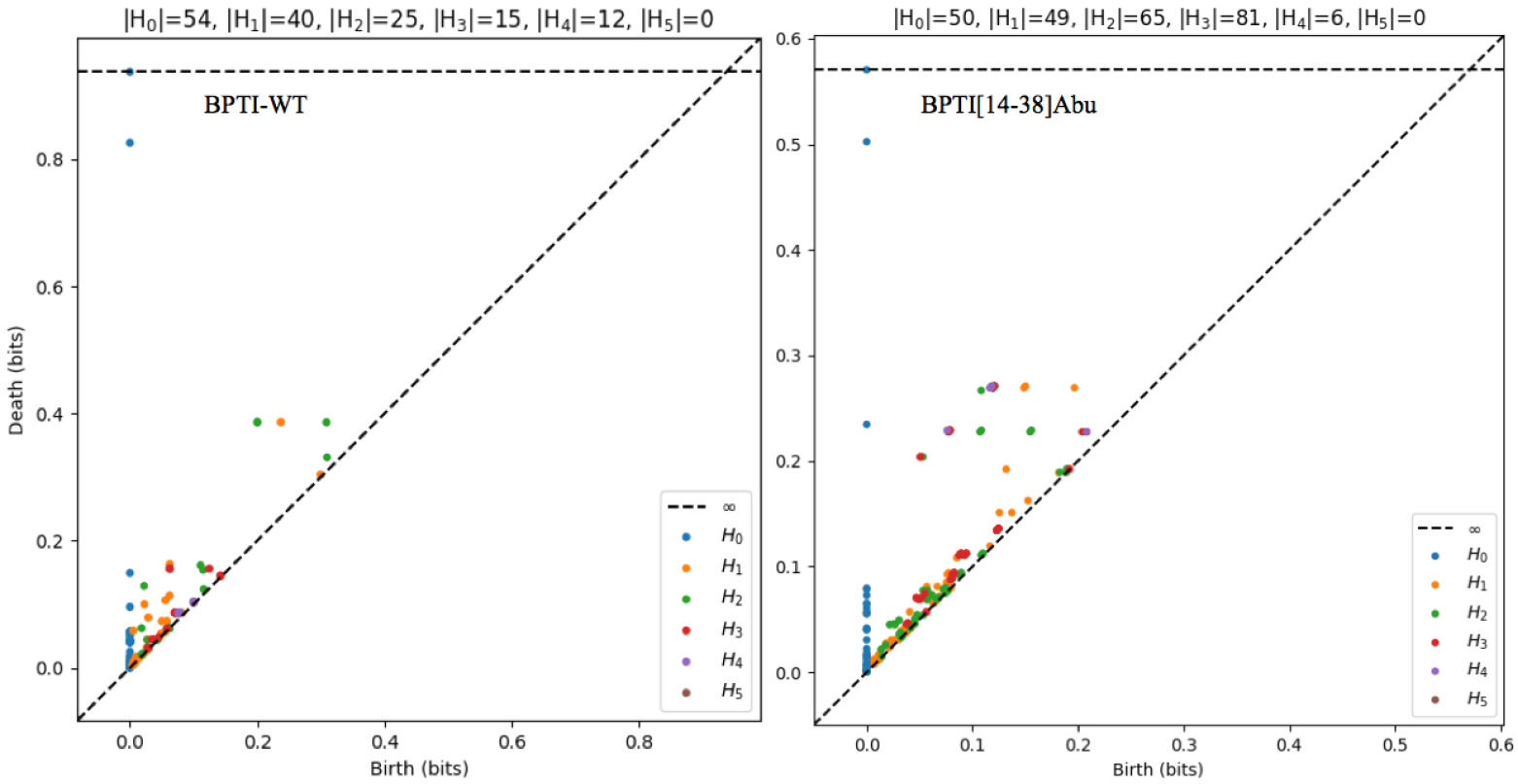
The topologies of the manifolds of BPTI wild-type and BPTI[14-38]Abu. The axes denote the length (in bits) at which a topological element appears (that is, the so-called birth) and the length at which it disappears (that is, the so-called death in bits). Here, *H*_0_ are isolated points (connected components), *H*_1_ line segments (holes), *H*_2_ triangles (cavities), *H*^*i*≥3^ poly-edges shapes (non-cavities, for example, *H*_3_ corresponds to a tetrahedron).

To quantify how far two topological spaces (ℳ_wt_, **g**) and (ℳ_mt_, **g**), given by the persistence diagrams in Fig. 8, we computed the Wasserstein and Bottleneck metric distances. Here, ℳ_wt_ = {*H*_0_, *H*_1_, *H*_2_, …}_wt_ and ℳ_mt_ = {*H*_0_, *H*_1_, *H*_2_, …}_mt_ are sets of different shapes, corresponding to the persistence diagrams (as in Fig. 8), which represents the topological spaces of BPTI wild-type and BPTI[14-38]Abu structures, respectively. Note that a persistence diagram is constructed for each shape (*H*_*i*_), when computing the metric distances, and each diagram (DG^(*i*)^) is a set of points in the two-dimensional diagram (*b, d*) and the diagonal line corresponds to the points of line *δ* = (*b* + *d*)*/*2. (Note that metric distances for *H*_0_ do not have any practical importance and are often omitted from this analysis. Besides, the points of diagonal lines *δ* are omitted, as well.)

First step is construction of a subset 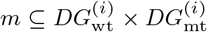 of so-called *matching* such that for every point 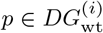 and 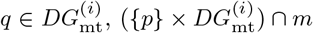 and 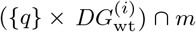 contain each a single pair. Then, the Bottleneck distance is defined as 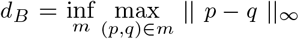 and the Wasserstein distance is defined as 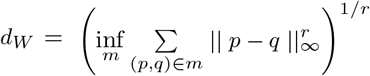 [18, 75] (and the references therein), for *r* ∈ [1, +*∞*).

In other words [22], the Bottleneck distance measures the largest distance of a pair of shape features *H*_*i*_ required to transform 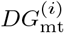 diagram into 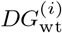; in contrast, the Wasserstein distance is the smallest of that distance. Importantly, while *d*_*W*_ distance is sensitive to the short-lived features (which are close to the diagonal line), the *d*_*B*_ distance is independent of them, and hence the Bottleneck distance does not count for the closeness of a small distance separated pair of points [26].

The computation results of the Bottleneck and Wasserstein distances are shown in Fig. 9 and Fig. 10, respectively, for transferring 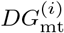 into 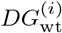 diagram (for *i* = 1, 2, 3, 4). In each graph, the green lines represent the matching set (*m*), and the length of a green line corresponds to the metric distance. Clearly, our results indicate that the number of significantly important features that match for wild-type protein (BPTI) and mutant protein (BPTI[14-38]Abu) reduces from *H*_1_ (holes) to *H*_4_ (non-cavities). That indicates that the protein structure topological space undergoes significant transitions in the dipole-dipole interaction network due to the mutations; in particular, dipole-dipole interaction networks of amino acids involving three or more residues become quite distant as common features for the wild-type and mutant protein. Therefore, interestingly, the topological space analysis should enable identification of the interaction network configuration space *transitions* due to the changes in the amino acid sequence.

**Figure 9.**
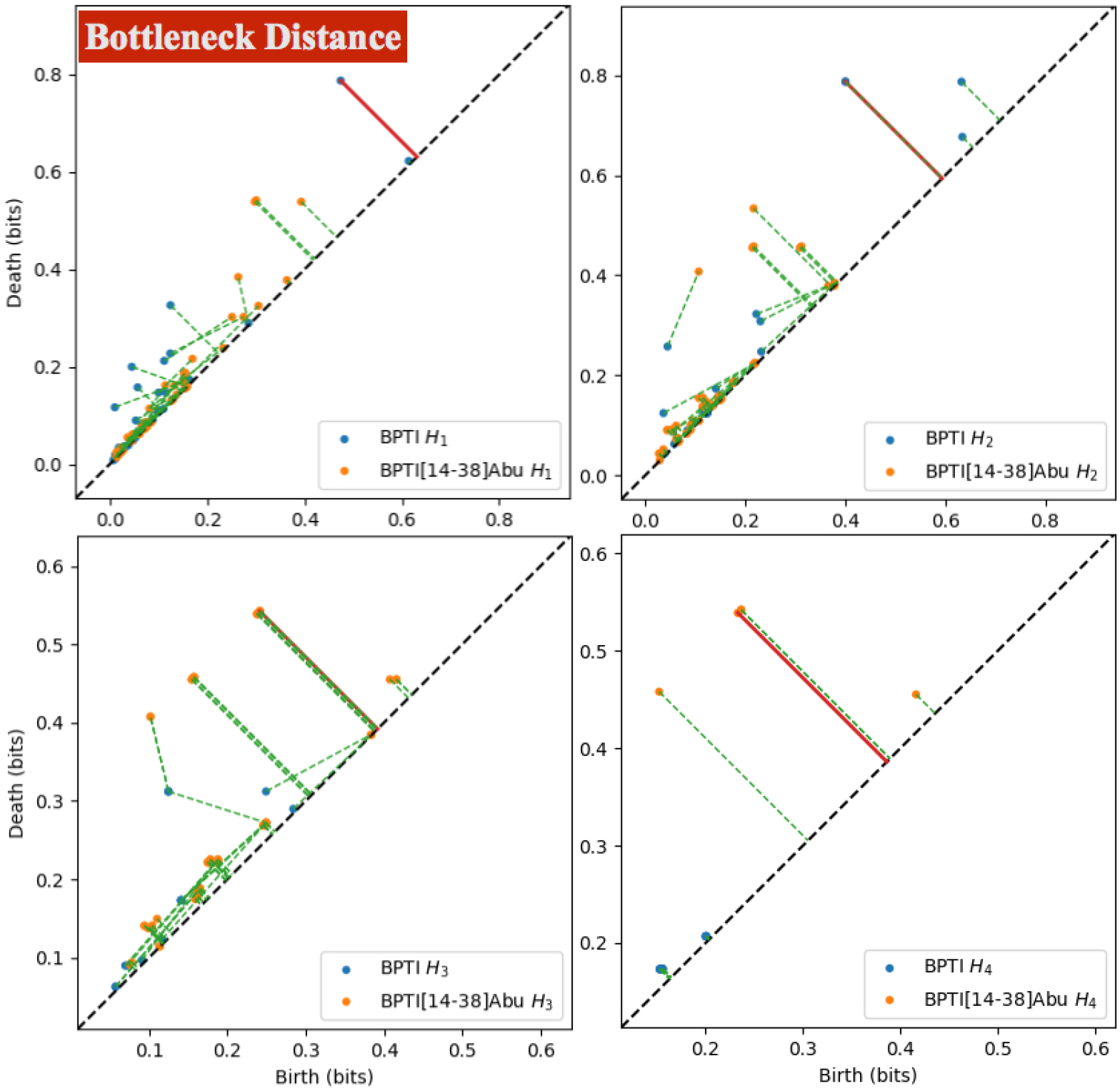
The Bottleneck distance metric between persistence diagrams of topological spaces for the embedded manifolds obtained for wild-type protein (BPTI) and mutant (BPTI[14-38]Abu). The green lines represent the matching set *m*, and their lengths depict the Bottleneck distances.

**Figure 10.**
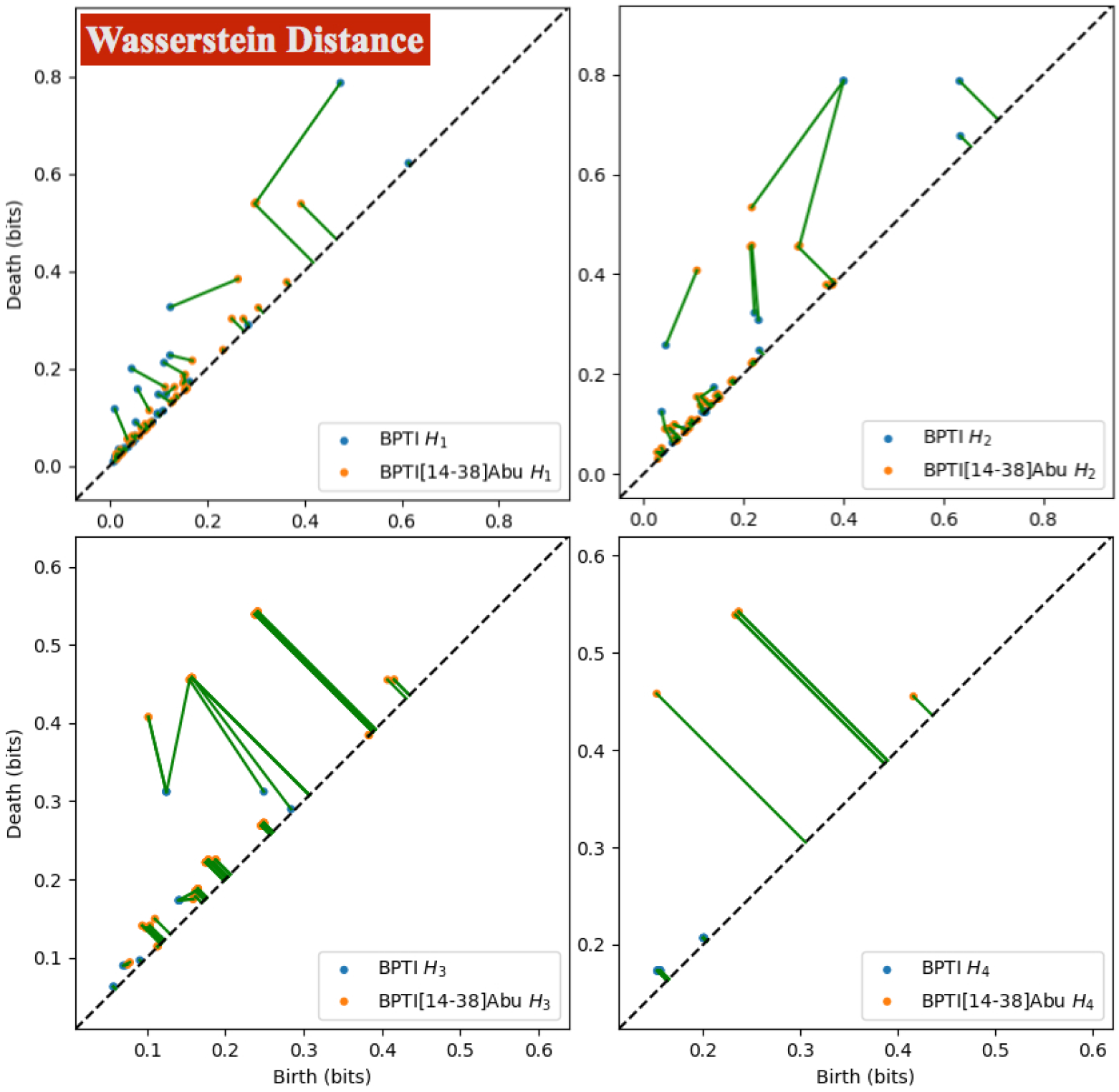
The Wasserstein distance metric between persistence diagrams of topological spaces for the embedded manifolds obtained for wild-type protein (BPTI) and mutant (BPTI[14-38]Abu). The green lines are elements of the matching set (*m*), and the length of a green line corresponds to the Wasserstein distance.

### 4.5 Embedded Manifolds of Asymmetric Kernel Information Flow

Next, we computed the embedding of the symmetric kernel matrix **K**^(*s*)^, presenting the adjacency matrix of a graph, into an *m*-dimensional space (typically, *m* = 15) and determined the embedded local coordinate vectors **x**_*u*_ for each vertex *u* (which represent the amino acids dipole positions of a protein structure). In addition, the spectral decomposition of the Laplacian matrix, based on the asymmetric kernel matrix **K** was used to compute the vector field components of the tangent vectors **v**_*u*_ (vectors of the local tangent space at **x**_*u*_ of the embedded manifold), characterising the embedded vector space of DiGraph with an adjacency matrix **K**.

Our results are presented graphically in Fig. 11. In the graphs, the first two components of the local coordinate vectors **x**_*u*_ (of a fifteen-dimensional embedded manifold, based in our computations) represent the points in a two-dimensional graph and the vectors depict the direction of the vector **v**_*u*_, representing the direction of the information flow in the embedded manifold. Our results indicate clearly distinct direction flows of the information for the wild-type BPTI-WT and BPTI[14-38]Abu structures; that is, there exist different dipole-dipole interaction graph topologies for the BPTI-WT wild-type and BPTI[14-38]Abu.

**Figure 11.**
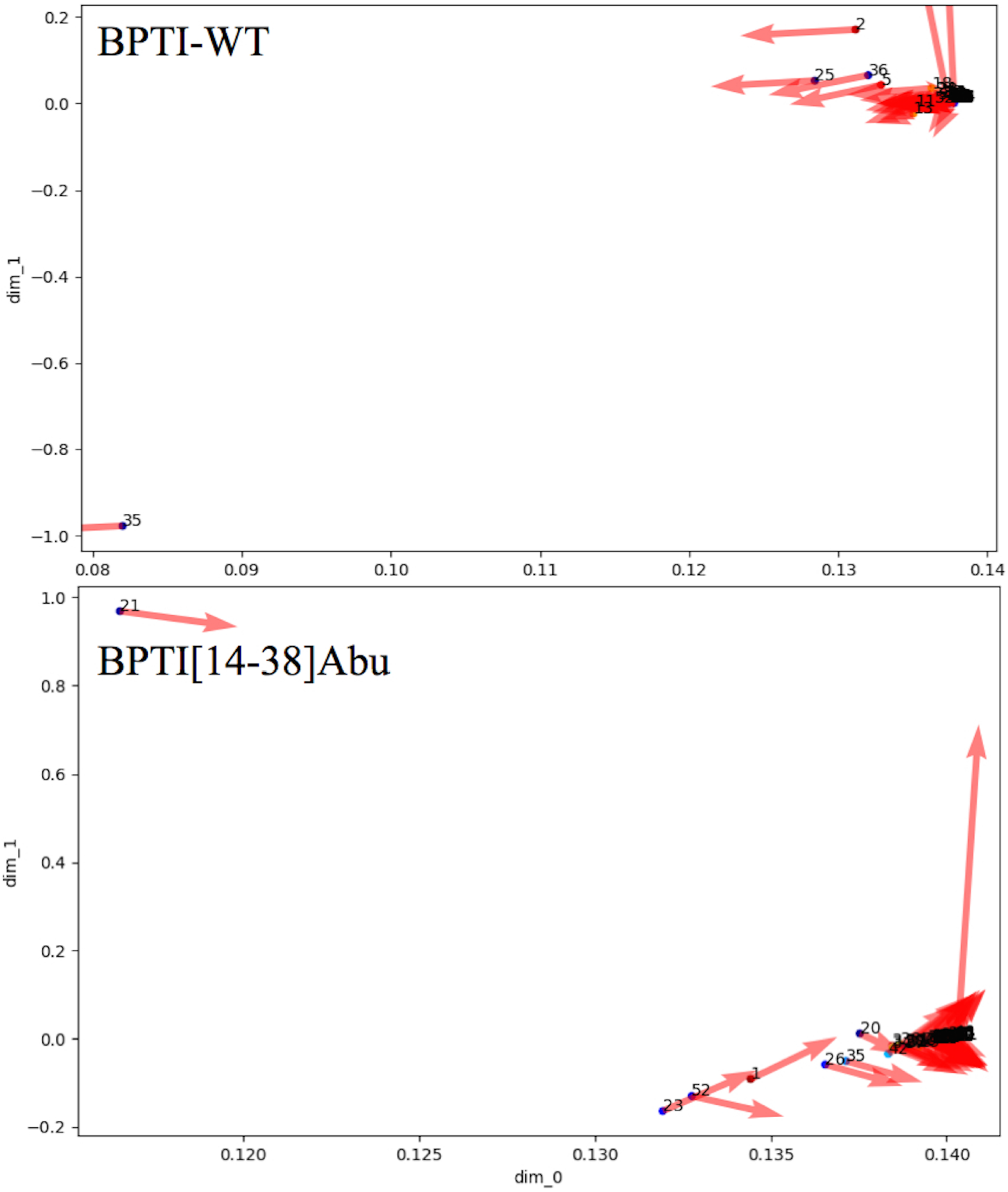
The embedded manifolds of BPTI wild-type and BPTI[14-38]Abu.

Furthermore, we performed the cluster analysis, based on the local coordinate vectors in the two-dimensional embedded manifold spaces, as shown in Fig. 12. Our data present different clusters formed for the wild-type protein BPTI-WT compared to the mutant analog BPTI[14-38]Abu. In addition, the direction of the information flow is clearly different for each protein structure within the different clusters.

**Figure 12.**
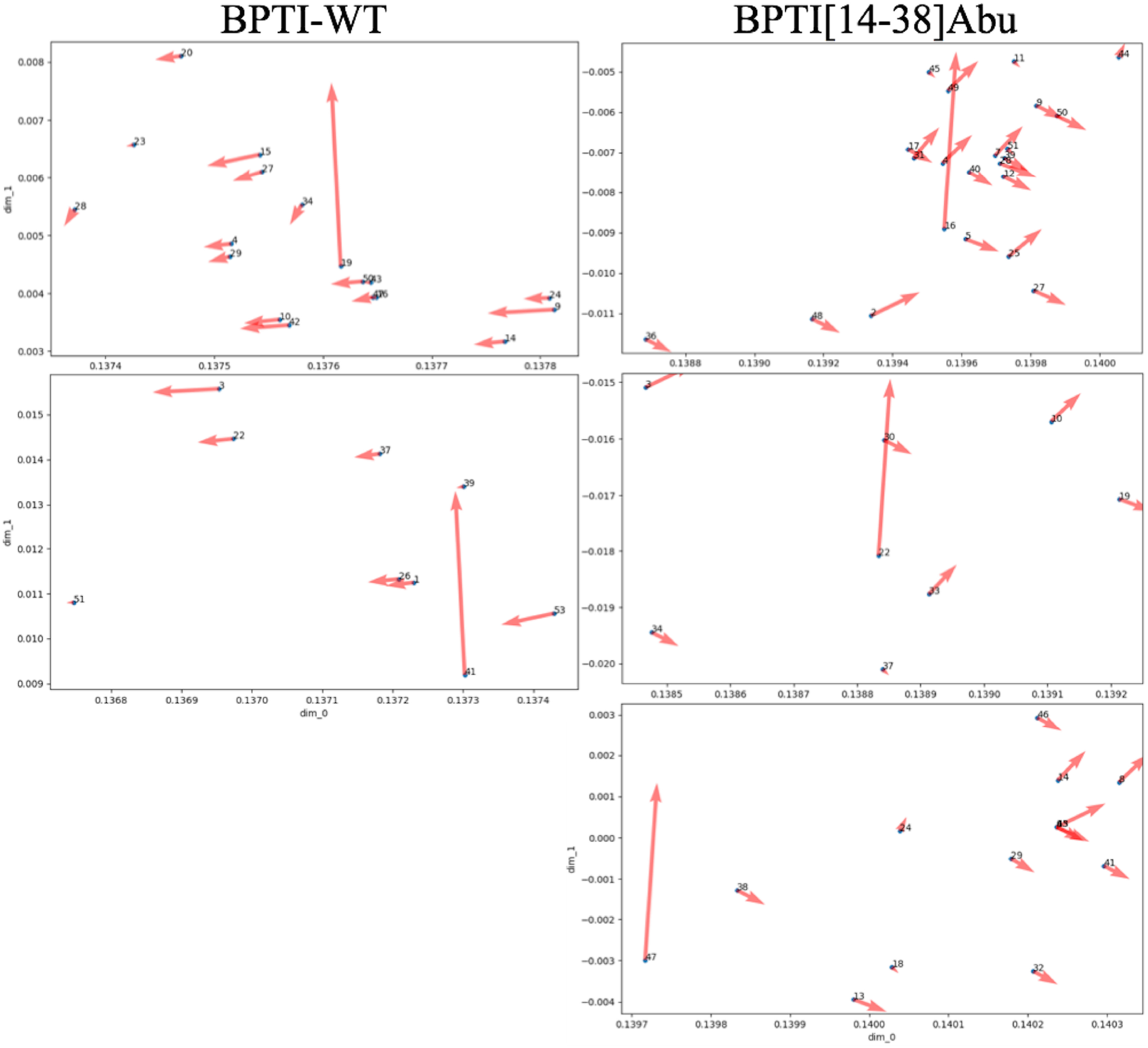
The embedded manifold clusters of dipole-dipole interactions for the BPTI wild-type and BPTI[14-38]Abu.

Importantly, these findings support the previously stated hypothesis that there exists a relationship between the topology space of the protein configuration manifold and configuration transitions associated with it. Furthermore, the topology can play a crucial role in studying the large-scale configuration transitions of different types of systems, which may highlight interesting phenomena, such as the folding pathways of protein structures. These results support the finding that the folding pathways of the BPTI-WT and BPTI[14-38]Abu are characterised by different collective dipole-dipole interactions of their configuration-embedded manifolds.

To further support our findings, we computed the curvatures of the embedded manifolds for each system. It is worth mentioning that the curvatures relate to the derivatives of the potential energy function *U*, representing a mapping of the topological space to a real-valued set. Furthermore, the changes in the curvatures at the local points of the embedded manifolds are associated with topological space changes of the protein configurations.

### 4.6 The Gaussian Curvatures of the Manifolds

Figure 13 presents the so-called Gaussian curvatures on the embedded manifolds for BPTI-WT and BPTI[14-38]Abu. The number of features used here is *m* = 15. The number of nearest neighbours was chosen according to *k* = int (0.8 · *N*_a.a._ + 0.5) where *N*_a.a._ is the number of amino acids in the protein structure. The Gaussian curvature is the product of the principal curvatures, as calculated using principal component analysis with a number of components equal to the number of embedded features: *ρ*_G_ = *k*_1_ · *k*_2_ where *k*_1_ and *k*_2_ are the first two largest eigenvalues [7].

**Figure 13.**
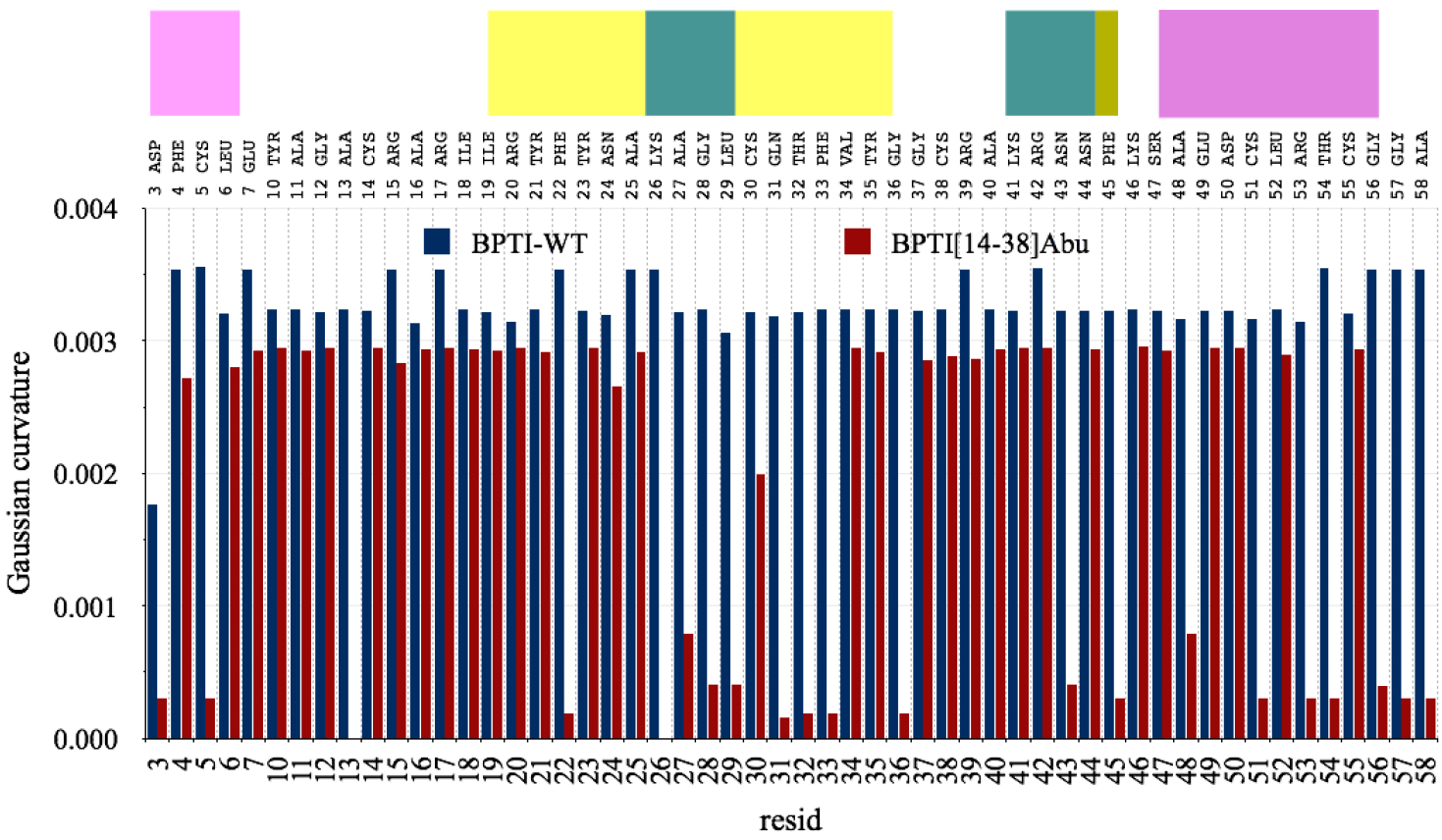
The Gaussian curvatures on the manifolds of BPTI wild-type and BPTI[14-38]Abu.

Figure 13 presents the Gaussian curvatures at every local coordinate point of a fifteen-dimensional embedded space of the protein structure for the BPTI wild-type and mutant analog. Interestingly, the Gaussian curvatures of wild-type protein structure BPTI-WT are, in general, greater than those of BPTI[14-38]Abu protein. In particular, in the region between amino acids 26 and 33, which represent the *β*_1_-sheet − turn − *β*_2_-sheet region, the values of *ρ*_G_ for BPTI[14-38]Abu are much smaller compared to *ρ*_G_ for wild-type BPTI. The same behaviour is observed for the amino acids Cys(Abu)5, Phe22, Gly36, Asn43, Phe45, Ala48, Cys(Abu)51, Arg53, and Thr54. That is most likely due to the long effective correlation times of the dipoles of these amino acids for the wild-type protein compared to the mutant protein.

A final note on the embedded manifold curvatures from the geometrical viewpoint. The larger the Gaussian curvature values, the smaller the curvature radii, and so more convex the hyper surface; on the other hand, small values of the Gaussian curvatures (and hence larger curvature radius) represent almost flat multidimensional spaces locally, which are supposed to optimise to the (local) minimum structure faster.

## 5 Conclusions

This study utilised the transfer entropy, based on the (*α, q*) framework, for measurement of higher-order correlations of the dipole-dipole interactions in proteins.

Furthermore, this study introduced topological data analysis to describe the directed graph embedded manifolds of the protein structure dynamics.

Moreover, the asymmetric kernel-directed graphs, determined by the transfer entropy, described the information flow in the embedded manifolds. Our primary goals were to characterise any changes in the topological spaces of the protein structures due to the mutations. In this study, we defined the embedded manifold of dimension *m* of the amino acid sequence interactions network using the so-called graph Laplacian matrix, which enabled us to determine the local embedded vector fields and coordinate vectors in this manifold for each amino acid, representing the vertices of either a directed or undirected graph.

We showed that encoding the amino acid sequence information in an *m*-dimensional embedded manifold is statistically efficient by allowing decoding of that information in a much lower-dimensional space. Besides, the topological data analysis enabled the observation of protein structure dynamics changes in a multi-dimensional manifold, for example, due to the amino acid mutations. Besides, in this study, we established a relationship between the changes in the topological spaces of embedded manifolds and the transitions in the configurational multilevel set potential energy function. In particular, the local Gaussian curvatures of the embedded manifolds explained those changes, supporting the finding that the information flows on the embedded manifolds are entropically driven. Moreover, the changes in the embedded manifolds determine the changes in the derivatives of the configurational entropy.

## Acknowledgments

The authors are grateful to the International Balkan University supporting the research activity.

## Data Availability

The supplementary materials used in this study are available within the article. The molecular dynamics simulations data used in this study are available from the corresponding author upon request.

## Footnotes

1 In Ref. [37], it was used as 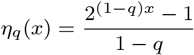. Our formula is slightly different because *H*_*α*_(*X*) is expressed in nats.

## Notes

### Competing Interest Statement

The authors have declared no competing interest.

